# Crystalline silica exposure induces multiple systemic autoimmune phenotypes including inflammatory arthritis and nephritis in Collaborative Cross mice with differing sub-clinical autoimmune profiles

**DOI:** 10.1101/2025.05.22.655650

**Authors:** Lisa M.F. Janssen, Caroline de Ocampo, Dwight H. Kono, Peter H.M. Hoet, K. Michael Pollard, Jessica M. Mayeux

## Abstract

Inhalation of crystalline silica dust, an occupational hazard, has been strongly associated with the development of autoimmune connective tissue diseases, such as rheumatoid arthritis and systemic lupus erythematosus. However, it remains unclear if silica-mediated autoimmune disease requires preexisting subclinical autoimmunity, and to what extent the severity of the preexisting condition influences silica-induced disease. The aim of the current study was to examine whether silica-mediated autoimmune disease requires preexisting subclinical autoimmunity, using the Collaborative Cross (CC) mouse model system, recognized for its ability to mimic the genetic diversity observed in human populations. Sixty-one CC strains were assessed for the presence of subclinical autoimmunity via autoantibodies and inflammatory markers. Six CC strains, chosen to represent a range of subclinical autoimmunity, were exposed transorally to 5 mg silica or PBS and examined 12 weeks later for lung inflammation, autoantibody responses, total immunoglobulin levels, and the manifestation of glomerulonephritis and autoimmune arthritis. Results indicated a spectrum of spontaneous subclinical autoimmunity among naive CC strains, with silica exposure leading to significant pulmonary inflammation and systemic autoimmunity, including glomerulonephritis and synovitis. Notably, strains with pre-existing subclinical autoimmunity showed more severe disease outcomes post-exposure.

## 1. Introduction

Autoimmunity is thought to result from a combination of genetics, environmental factors, and stochastic events [1–4]. Many autoimmune diseases are preceded by the presence of immune dysregulation and clinical manifestations that may be present years before a clinical diagnosis is made [5, 6]. This phase of autoimmune disease development has been variously labelled as sub-clinical, pre-clinical, asymptomatic, or incomplete autoimmunity [2, 6–8] and is characterized by the presence of autoantibodies and changes in other immune biomarkers including cytokines and chemokines [5, 6]. Autoantibodies are hallmarks of autoimmunity and can be important in diagnosis and disease severity [9–12]. Disease relevant autoantibodies can predate, by up to a decade or more, the onset of clinical symptoms of systemic lupus erythematosus (SLE) [13], Sjögren’s syndrome (SS) [14], rheumatoid arthritis (RA) [15–17], and systemic sclerosis (SSc) [18]. Importantly, other parameters of sub-clinical autoimmunity, such as proinflammatory cytokines and chemokines, when coupled with autoantibody levels, can improve prognostic precision [19]. Events that trigger sub-clinical autoimmunity are not clearly understood but genetics and environmental exposures are important risk factors [5, 6, 8]. Although epidemiological studies confidently link environmental factors to a number of systemic autoimmune diseases [20–23], it is not clear if the resulting disease is influenced by preexisting sub-clinical autoimmunity, is a specific response mediated by the exposure, or a combination of both susceptibility to autoimmunity and specific environmental factors [24–27].

Occupational crystalline silica dust exposure is a well-established risk factor for autoimmune connective tissue diseases (CTDs) in humans, including SLE, RA, and SSc [23, 28–33]. Pulmonary exposure to silica dust is a significant event in the tissue damage and chronic inflammation that precedes development of both silicosis and features of autoimmunity [2, 34]. These events are consistent with the hypothesis that autoimmune biomarkers at mucosal sites are important indicators of immune processes necessary for development of autoimmune diseases [8, 35], and therefore implicate the lung as an important site in the origin of CTDs linked to silica dust exposure [2]. While silica exposure can significantly increase the risk of developing a specific CTD [32, 36], the development of immunological features consistent with sub-clinical autoimmunity is a more common finding. Proinflammatory cytokines and lung inflammation, precursors to silicosis [37], are observed in 47–77% of individuals following silica exposure [38]. Among silicosis patients, over 65% may develop hypergammaglobulinemia [39], and the prevalence of anti-nuclear autoantibodies (ANA) can reach 34% or more [40, 41]. But only a small proportion of exposed individuals will progress to autoimmune disease specific pathology [2, 28, 31]. These findings are consistent with a phenotypic progression that begins with sub-clinical features including activation of proinflammatory cytokine production, inflammation of the lung leading to activation of adaptive immunity, breaking of tolerance, production of autoantibodies, and, in susceptible individuals, development of pathological disease [33, 37, 42–45]. Experimental model studies using both inbred [46] and outbred [34] mouse strains, including those that develop systemic autoimmunity [24, 47], support these observations. However, little is known about the contribution that sub-clinical autoimmunity plays in the development and severity of autoimmune diseases linked with exposure to silica.

Numerous animal models have been used to study idiopathic and induced systemic autoimmunity, particularly SLE [48, 49]. Animal model studies of idiopathic systemic autoimmunity have been augmented by the generation of recombinant inbred (RI) strains having various levels of severity of the parental autoimmune phenotype [50–53]. However, RI strains based on severe parental autoimmune phenotypes are not ideal for studies of the progression from non-autoimmune, to sub-clinical, and to systemic autoimmune disease. This limitation has been overcome by generation of a large panel of multi-parental RI strains, called the Collaborative Cross (CC) [54, 55]. Based on five laboratory strains (A/J, C57BL/6, 129S1/SvlmJ, NOD/ShiLtJ, NZO/HiLtJ) and three wild-derived strains (WSB/EiJ, PWK/PhJ, CAST/EiJ), the CC RI panel captures over 90% of the worldwide genetic diversity in mice. The panel is a unique mammalian resource with genome-wide genetic variation randomized across a large, heterogeneous and reproducible population [55, 56]. Thus, CC RI strains exhibit heritable variations of immune phenotypes [57–60] including a spontaneous inflammatory disease resembling colitis in the CC011 strain [61]. While evidence of sub-clinical autoimmune manifestations have not been formally described in CC RI strains, the related Diversity Outbred (DO) mice, a genetically heterogeneous stock derived from the same eight founder strains, spontaneously develop several different autoantibodies and a significant percentage are susceptible to the development of autoimmune pathology following silica exposure [34]. Thus, CC RI strains may constitute a powerful genetically heterogeneous resource for studying environmental factors that may regulate the initiation and progression of autoimmunity from sub-clinical to clinical systemic disease.

In this study, we tested the hypothesis that the genetic heterogeneity of naïve CC RI mice results in a spectrum of sub-clinical autoimmunity that can be used to examine how these early, non-diagnostic, features of autoimmunity impact the development and severity of silica-induced systemic autoimmune diseases. Serological analysis of 61 naïve CC RI strains identified a range of sub-clinical ANA and proinflammatory cytokine and chemokine responses. Using this information, six strains, representing varying levels of sub-clinical autoimmunity, were exposed to crystalline silica. The resulting responses in pulmonary inflammation, cytokine and chemokine expression, and autoantibody profiles supported the proposition that pre-existing susceptibility to sub-clinical autoimmunity can be exacerbated by silica exposure leading to more severe pathology strikingly demonstrated by development of inflammatory arthritis and lupus nephritis. These findings argue that although CC RI strains are based on the same eight founder strains [56], their genetic heterogeneity results in a spectrum of sub-clinical autoimmunity that can result in diverse pathological responses to crystalline silica exposure that mimic those found in occupationally exposed humans. CC RI mice thus constitute a valuable resource to study how subtle genetic differences affect the role of sub-clinical autoimmune phenotypes in the development and severity of human autoimmune diseases linked to occupational and environmental exposures.

## 2. Materials and methods

### 2.1. Serum samples from naïve CC mice

Serum samples from male and female Collaborative Cross (CC) Recombinant Inbred (RI) breeders were obtained from the Systems Genetics Core Facility at the University of North Carolina at Chapel Hill. Samples were obtained from strains maintained under Specific Pathogen Free (SPF) conditions (CC1 cohort) or under ultra-clean Barrier conditions (CC2 cohort). Samples were available from 61 strains totaling 392 mice (6-10 mice/strain). The age range was 12.3 – 64.7 weeks, with an average of 32 weeks; the average age was 32.2 weeks for females and 31.9 weeks for males.

### 2.2. Breeding and maintenance of CC strains

Male and female mice from six CC RI strains (CC081/Unc, CC061/GeniUnc, CC041/TauUnc, CC039/Unc, CC072/TauUnc, CC046/Unc) were obtained from the Systems Genetics Core Facility at the University of North Carolina at Chapel Hill. All mice were bred and maintained under SPF conditions. Animal rooms were kept at 20-22°C and 60-70% humidity, with a 12 hr/12 hr light-dark cycle, and sterilized cages were replaced each week with fresh water and food (autoclaved standard grain diet 7012, Teklad, Envigo, Madison, WI, USA) to which the mice had access ad libitum.

### 2.3. Exposure to crystalline silica

Crystalline silica (SiO_2_) (Min-U-Sil-5, average particle size 1.5-2 μm; U.S. Silica Company, Frederick, MD) was prepared and reconstituted in PBS as previously described [62]. Immediately prior to use, silica was disbursed by sonication. Male and female mice (8-11 weeks old) were exposed to a single dose of 5 mg by transoral (TO) instillation in a volume of 50 µl in PBS as described previously [63, 64]. The dose of 5 mg was chosen based on calculations of human lifetime exposure to respirable crystalline silica and an equivalent exposure for mice, and represent approximately 60% of a human lifetime exposure at the recommended NIOSH exposure limit of 0.05mg/m^3^/d [24]. An instillation volume of 50 µl was used as this results in the most consistent lung inflammation [34, 65]. The experimental protocol was approved by the Institutional Animal Care and Use Committee at The Scripps Research Institute (protocol# 08-0150), and use of silica was approved by the TSRI Department of Environmental Health and Safety.

### 2.4. Induction of Collagen Antibody Induced Arthritis (CAIA)

CAIA was induced in C57BL/6 mice by subcutaneous injection of 1.5 mg of a cocktail of 5 mouse anti-type II collagen monoclonal antibodies (10mg/ml) followed by 50 ugs of LPS on day 3 (Chondrex Inc, Woodinville, WA. catalog #53040). Paw inflammation was evaluated on days 0, 2, 7, 10, 14 as described by the manufacturer, and synovitis and bone erosions by ankle joint histopathology on day 14 as described below in section 2.6.

### 2.5. Lung and kidney histopathology

Lungs, tracheobronchial lymph node (TBLN) and kidneys were excised and fixed for 48 hours in zinc formalin before paraffin embedding and sectioning (4um). Lung and TBLN sections were stained with Hematoxylin and Eosin (H&E) and kidney sections stained with Periodic Acid–Schiff (PAS). Embedding, sectioning and staining were performed by the Microscopy and Histology Core Facilities of the La Jolla Institute for Immunology (LJI). Slides were scanned using a Leica AT2 whole slide scanner (Microscopy Core, Scripps Research) and images stored on Digital Image Hub (Slidepath, Dublin, Ireland). Lung inflammation was assessed, under blinded conditions, by determining the percent of each of the 5 lobes affected by alveolitis (increased number of macrophages, accumulations of inflammatory cells and thickening of alveolar wall) or peribronchitis/perivasculitis (accumulations of inflammatory cells around the vasculature and bronchioles). Alveolitis and peribronchitis/perivasculitis were scored from 0-500 and the two values combined to give a total lung score (TLS) out of 1,000 [34]. Inflammation and silicosis were further characterized by scoring for lung granuloma (0-3) (formation of silica-containing necrotic areas, surrounded by inflammatory cells), proteinosis (-or +), and fibrosis (0-3). Fibrosis was scored based on H&E-stained sections, and confirmed by Pico Sirius Red and Masson’s Trichrome staining. TBLN involvement was assessed by taking the TBLN weight, and scoring for the formation of silicotic nodules in the TBLN. Glomerulonephritis was scored, under blinded conditions, on a 0-4 scale under blinded conditions, as described by Koh et al. [66].

### 2.6. Ankle joint histopathology

Ankle bones were collected, placed in cassettes and fixed by stirring for 72 hours in zinc formalin and then decalcified using 5% formic acid. Decalcification, preparation of paraffin sections (4 μm), and H&E staining was performed by the Microscopy and Histology Core Facilities of LJI. Stained slides were scanned using a Leica AT2 whole slide scanner (Microscopy Core, Scripps Research) and images stored on Digital Image Hub (Slidepath, Dublin, Ireland). Arthritis was scored for synovitis and bone erosion, both on a scale from 0 – 3, under blinded conditions, as described by Hayer et al. [67].

### 2.7. Immunofluorescent staining of kidney sections

For detection of renal deposits of IgG and C3, tissue was embedded in OCT Tissue-Tek® (Sakura Fintek, USA), frozen and 6 um sections cut using a cryostat. Slides were incubated with ice cold acetone (100%, 10 min), washed in PBS, and blocked in 2% BSA for 45 min. before overnight incubation at 4°C with an antibody-cocktail containing AF555-conjugated-anti-IgG (Southern Biotech, Catalog #: 1030-32) diluted 1/50 and FITC-conjugated anti-C3 (ICL Lab, Catalog#: GC3-90F-Z) diluted 1/100. Slides were then washed with PBS, and coverslips mounted using Vectashield anti-fade mounting medium with DAPI (Vector Laboratories Inc., Ref. H-1200). Sections were visualized using a Keyence BZ-X710 fluorescence microscope, and digital images containing up to 5 glomeruli were captured using its CCD camera.

### 2.8. Antinuclear antibodies (ANA) using Indirect Immunofluorescence (IIF)

Serum ANA were detected as previously described [68] using a 1:40 (naïve CC mice) or 1:100 (silica study) dilution of serum on Hep2 ANA slides (Innova Diagnostics, San Diego, CA). Bound antibody was detected with goat anti-mouse IgG Alexa Fluor 488 (Life Technologies, Carlsbad, CA). Slides were mounted using Vectashield Mounting Medium (Vectorlabs, Burlingame, CA), and visualized using an Olympus BH2 microscope (Olympus America Inc, Cypress, CA). Digital images were captured using a LEICA DFC 365 FX camera and analyzed using Leica Application Suite AF software (LEICA Microsystems, Buffalo Grove, IL). ANA was assessed based on fluorescence intensity on a scale of 0-4, with a score of 1 or higher considered positive [69].

### 2.9. ELISA for serum immunoglobulins and autoantibodies

Quantitative measurement of antibodies against ANA, ENA6 (Sm, -RNP, -SS-A (60kDa and 52kDa), - SS-B, Jo-1 and -Scl-70), CCP3 (Cyclic Citrullinated Peptide, 3rd generation), and chromatin were determined by ELISA (Inova Diagnostics, San Diego, CA) modified to detect murine samples using a goat-anti mouse IgG HRP antibody (Invitrogen #34137). Determination of antibody units was achieved using human negative, low positive, and high positive controls as described by the manufacturer (Innova Diagnostics) [34]. The units for each sample were calculated by dividing the average OD of the sample by the average OD of the human Low Positive. The result was multiplied by the number of units assigned to the human Low Positive (25 Units). IgM Rheumatoid Factor (RF) was measured using a sandwich ELISA, performed by coating IgG1κ (BD Pharmingen, San Diego, CA, USA) at 1 μg/ml onto NUNC plates (Thermo Fisher Scientific) and blocking with PBS containing 1 mg gelatin/ml. Serum was diluted 1:100, and bound RF detected with goat anti-mouse IgM HRP (Invitrogen, Carlsbad, CA, USA) at a 1:1,000 dilution. Serum from C57BL/6J-Lpr and NOD/ShiLtJ mice was used as positive controls for reactivity of mouse sera for all ELISAs and used for inter-plate calibration.

### 2.10. High-Throughput Protein Microarray autoantibody Profiling

High-throughput profiling of IgG autoantibodies in serum against a broad range of autoantigens (AAgs) was performed by the Microarray and Immune Phenotyping Core Facility at The University of Texas Southwestern Medical Center. Briefly, serum samples were pre-treated with DNAse I to remove free-DNA, and subsequently diluted at 1:50 and hybridized to protein array plates coated with 120 antigens and 6 controls. The list of included antigens can be found in Supplementary Table 1. The antibodies binding to the antigens on the plates were detected with Cy3-conjugated anti-mouse IgG or IgM (1:2000, Jackson ImmunoResearch Laboratories, PA). Fluorescent images were captured with a Genepix 4200A scanner (Molecular Devices, CA) and transformed to signal intensity values using GenePix 7.0 software. Images were background-subtracted and normalized to internal controls for IgG. The processed signal intensity value for each autoantibody was reported as antibody score (Ab-score), which is expressed based on the normalized signal intensity and signal-to-noise ratio (SNR) using the formula: Ab−score=log2(NSI∗SNR+1). Normalized signal intensity (NSI) values were represented in a heatmap, organized by unsupervised hierarchical clustering generated using ClustVis [70]. Significant differences between PBS- and c-silica-exposed mice were tested using t-tests with compensation for multiple testing.

### 2.11. Inflammatory mediators

Sera were tested for inflammatory markers (Rantes/Ccl5, M-CSF, G-CSF, IFN-γ, MIG/Cxcl9, IL-2, IL-5, IL-6, IL-17A, MIP-1β/Ccl4) by MULTIPLEX Map assay according to the manufacturer’s instructions (Millipore Sigma, Burlington, MA). Markers were chosen based on human autoimmune disease biomarkers argued to distinguish between healthy ANA negative individuals (Rantes/Ccl5 and M-CSF), patients with the autoimmune disease SLE that were ANA positive (G-CSF, IFN-γ, MIG/Cxcl9, IL-2, IL-5, IL-6, IL-17A), and healthy ANA positive individuals (MIP-1β/Ccl4) [9, 19]. For each marker, a mean for all strains was determined and a relative change from the strain mean was calculated for each individual strain. Strains that had higher mean levels of inflammatory markers have a positive relative change to the mean for all strains while strains that had lower mean levels of inflammatory markers have a negative relative change to the mean for all strains.

### 2.12. Determination of the presence and absence of sub-clinical autoimmunity

Data from autoantibody and inflammatory marker analysis of serum from naïve CC RI breeders (obtained from the Systems Genetics Core Facility, UNC, Chapel Hill) were used to identify those CC RI strains exhibiting serological features of sub-clinical autoimmunity. To decide which strains were considered positive for ANA, a cutoff point was determined using the mixtools package in R to identify two normal distributions (negative and positive ANA) within a bimodal population in the ANA ELISA dataset. Hierarchical cluster analysis for inflammatory mediators was done using Cluster 3.0 and Java TreeView using a relative change value over the mean of all included strains for each phenotype and Pearson correlation (uncentered) similarity metric as described [34] to identify two major groups (negative and positive inflammatory marker) using the relative change compared to the strain mean for each marker.

### 2.13. Statistical analysis

Data are expressed as mean ± SEM unless otherwise stated. Statistical analyses were performed using GraphPad Software V6 (San Diego, CA). Comparisons were performed using Two-Way ANOVA with Šídák’s correction for multiple testing. p<0.05 was considered significant. P values were further indicated as follows: *p<0.05, **p<0.01, ***p<0.001, ****p<0.0001.

## 3. Results

### 3.1. Collaborative Cross RI strains manifest varying degrees of subclinical autoimmunity

To assess the presence of sub-clinical autoimmunity in naïve CC RI mice, serum samples from 61 strains were tested for ANA and 10 inflammatory markers (Figure 1). CC RI strains showed a wide range of ANA responses, from negative to highly positive. Shown in Figure 1A are the 32 strains that had a mean ANA value above the positive cutoff point. Notably, most strains with a positive mean ANA score contained 50% or more ANA-positive mice (26/32, 81%). Analysis of individual strains revealed no correlation of age and/or sex with ANA values (data not shown). Inflammatory marker analysis using hierarchical clustering identified two major groups, positive (19/61, 31%) and negative (42/61, 69%), as well as a number of subgroups within both the positive and negative groups (Figure 1B).

**Figure 1:**
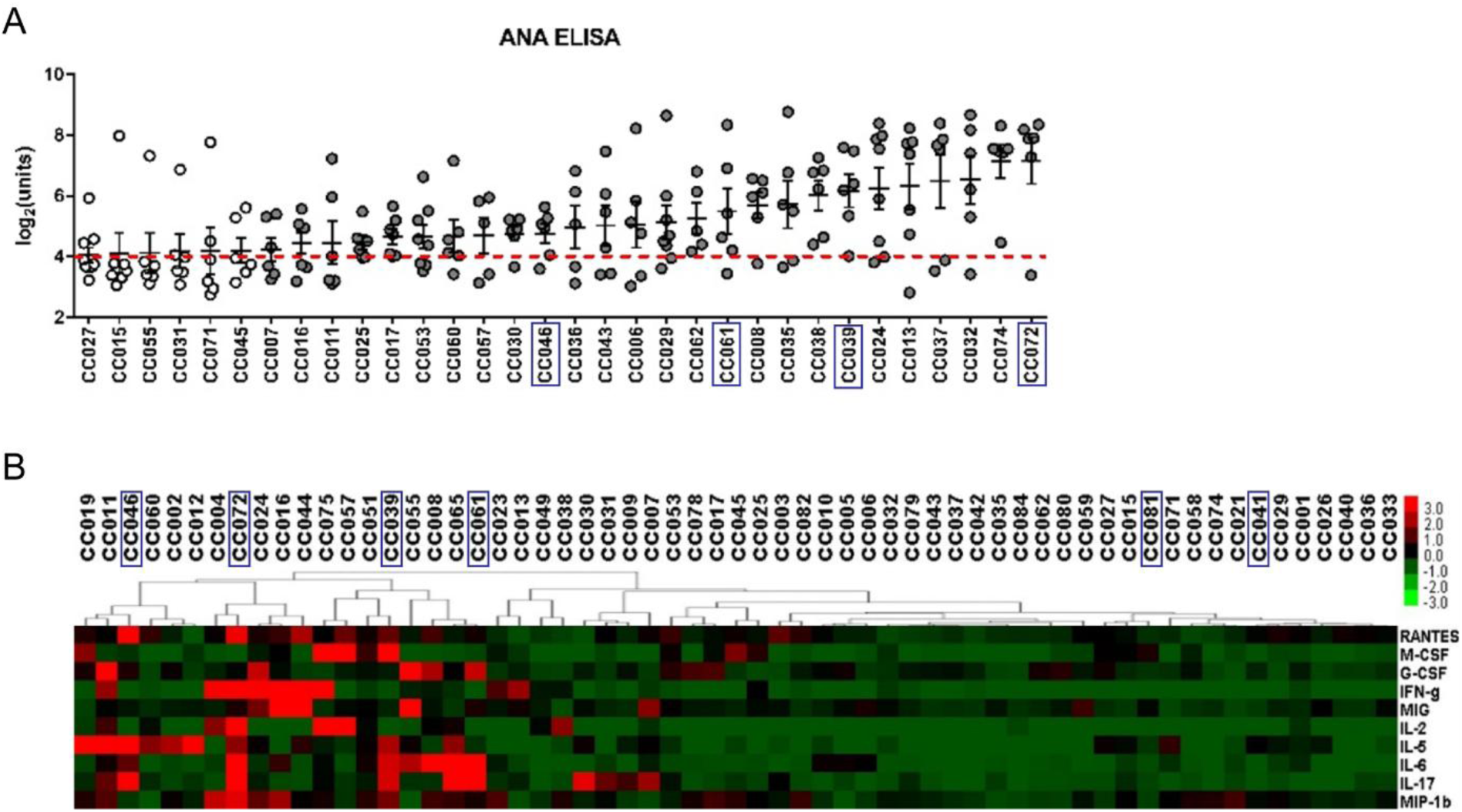
Subclinical autoimmunity in 61 Collaborative Cross RI strains. (A) ANA in 61 CC RI strains. Sera from 61 CC RI strains were tested for IgG ANA by ELISA. Shown is the subset of 32 strains with a mean score above the positive cutoff value (red dotted line). Strains with ≥50% ANA+ mice are shown in grey, and strains with <50% ANA+ mice are in white. Strains selected for silica exposure are shown boxed. N = 6-10 mice/strain. (B) Hierarchical cluster analysis of inflammatory markers in CC RI mice. Sera from 61 CC strains were tested for the presence of Rantes/Ccl5, M-CSF, G-CSF, IFN-γ, MIG/Cxcl9, IL-2, IL-5, IL-6, IL-17A, MIP-1β/Ccl4 by MULTIPLEX Map assay (Millipore Sigma, Burlington, MA). Hierarchical Cluster Analysis was done using Cluster 3.0 and Java TreeView using a relative change value for each phenotype and Pearson correlation (uncentered) similarity metric. Strains selected for silica exposure are shown boxed. N = 6-10 mice/strain.

### 3.2. Selection of CC RI strains to study silica-induced autoimmune disease

To study the role of sub-clinical autoimmunity in the development of silica-induced systemic autoimmune disease, six CC RI strains were selected based on their level of subclinical autoimmunity, spanning a range from negative for ANA and inflammatory markers, to positive for both ANA and inflammatory markers as shown in Table 1 and in boxes on Figure 1A and 1B. Two ANA and inflammatory marker negative strains were chosen, and four ANA and inflammatory marker positive.

**Table 1:**
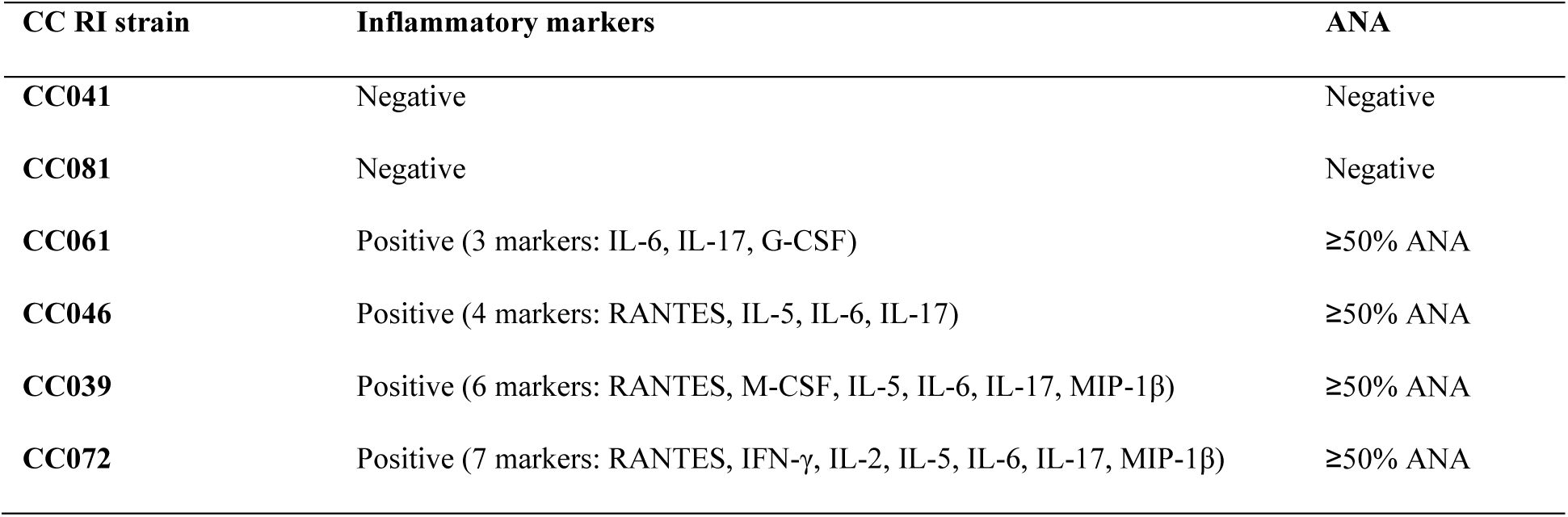
Subclinical autoimmunity (ANA and inflammatory markers) in CC RI strains selected for silica exposure.

Figure 2 shows the study design used to evaluate the extent of silica-induced disease, including lung inflammation, humoral autoimmunity, and autoimmune disease manifestations, in the selected CC strains 12 or 20 weeks after transoral exposure to crystalline silica.

**Figure 2:**
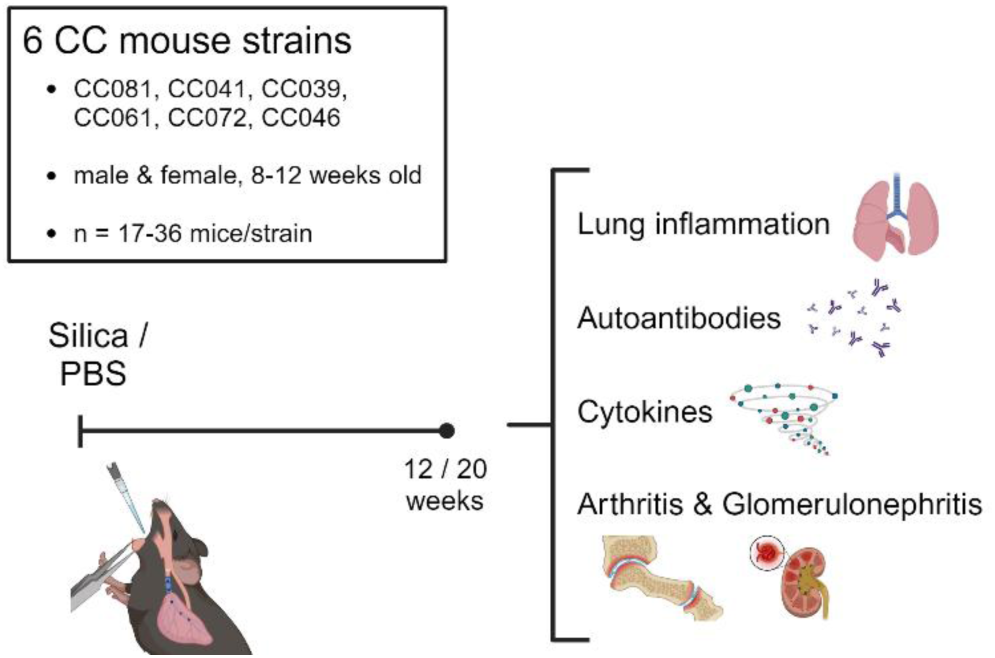
Study design. Male and female mice in age- and gender-matched groups were exposed to either silica (5 mg in 50 µl PBS) or 50 µl PBS via transoral instillation, and harvested 12 or 20 weeks post-exposure.

### 3.3. Silica exposure mediates severe inflammatory arthritis in CC039 mice

Strikingly, CC039 mice developed severe inflammatory arthritis upon silica exposure. Arthritis was manifested as synovitis by 12 weeks (Figure 3A) with characteristic thickening of the synovium and infiltrations of inflammatory cells, mainly neutrophils, which became pronounced by 20 weeks. Synovitis was exclusively observed in female mice, at 12 or 20 weeks post-exposure (Figure 3B). Bone erosions were observed by 20 weeks post-exposure (Figure 3C). The inflammatory arthritis observed in CC039 mice resembled that seen in other models of inflammatory arthritis, such as collagen-antibody induced arthritis (Figure 3D), albeit less severe (Figure 3E). None of the other silica exposed CC RI strains showed any signs of arthritis post-silica exposure (data not shown), nor did C57BL/6J mice (Figure 3C).

**Figure 3:**
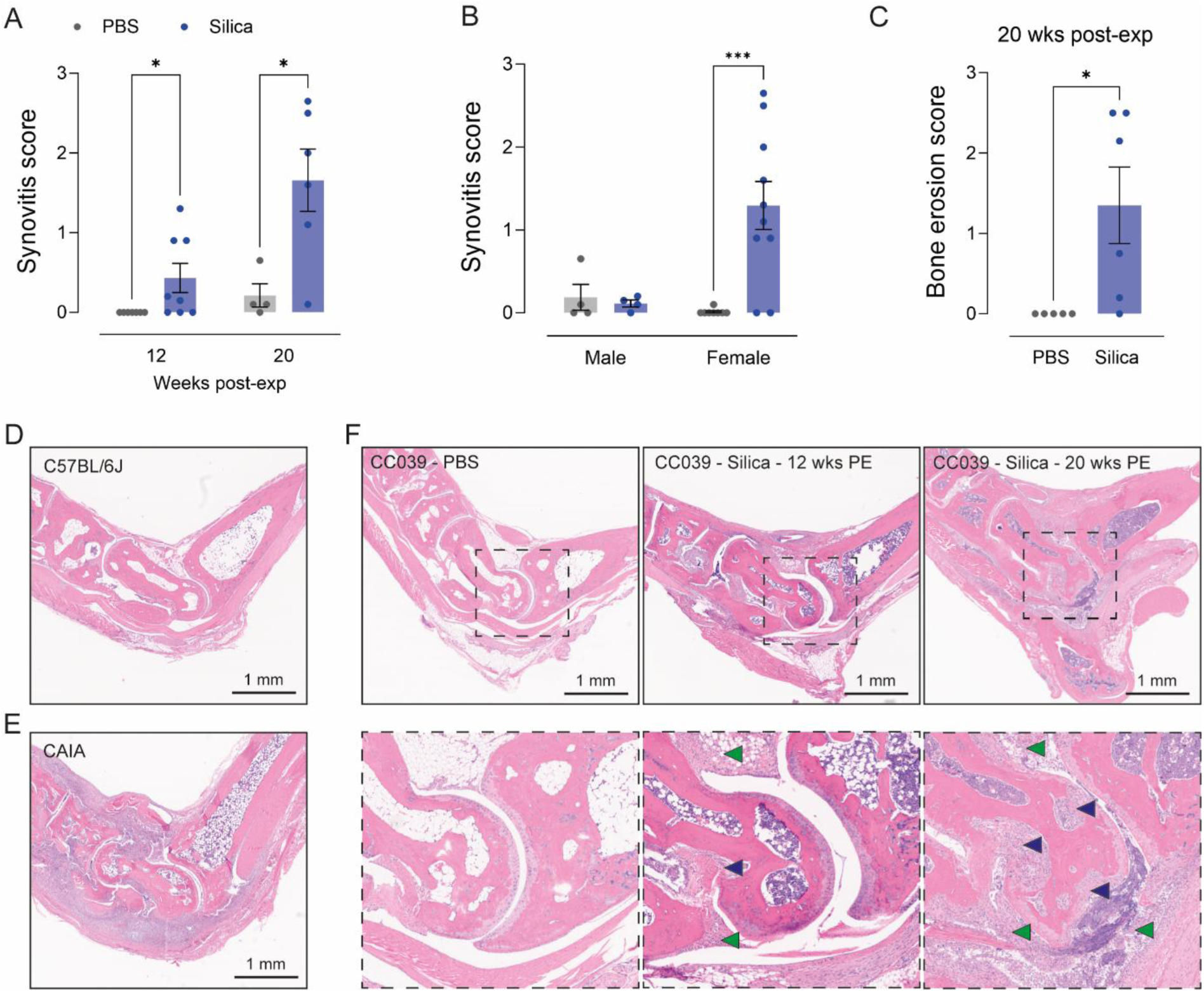
Silica exposure mediates severe arthritis in CC039 mice. (A) Synovitis scores of CC039 mice, 12 and 20 weeks post-exposure. (B) Comparison of synovitis scores (12 and 20 week cohorts combined) between male and female mice. (C) Bone erosion scores of CC039 mice, 20 weeks post-exposure. Comparisons were performed using Two-Way Anova with Šídák’s correction for multiple testing (A&B) or a T-test (C). *p<0.05, **p<0.01, ***p<0.001, ****p<0.0001. (D) H&E-stained ankle joint of a C57BL/6J mouse as a negative control. (E) H&E-stained ankle joint of a collagen-antibody induced arthritis (CAIA) model as a positive control. (F) H&E-stained ankle joints of CC039 mice, PBS-exposed, silica-exposed 12 weeks post-exposure and silica-exposed 20 weeks post-exposure. Bone erosion is indicated with blue arrows, and synovitis with green arrows.

### 3.4. Silica exposure exacerbates glomerulonephritis in CC061 mice

Silica exposure in CC061 mice resulted in a lupus-like phenotype including development of glomerulonephritis (GN) to a significantly greater extent compared to PBS mice (Figure 4A). A significant sex effect was observed, with more GN in female PBS- and silica-exposed CC061 mice (p = 0.0158 for sex effect) (Figure 4B). However, the lack of an interaction effect shows that both male and female mice are impacted by silica exposure to a similar extent. GN manifested as enlarged glomeruli, proliferation and glomerular infiltration (Figure 4C). There were no signs of lymphocytic infiltration in the kidneys. Immunofluorescent staining showed immune-complex deposits of IgG and complement factor 3 (C3) in the glomeruli of mice with histological evidence of GN (Figure 4D). None of the other silica-exposed CC strains developed GN.

**Figure 4:**
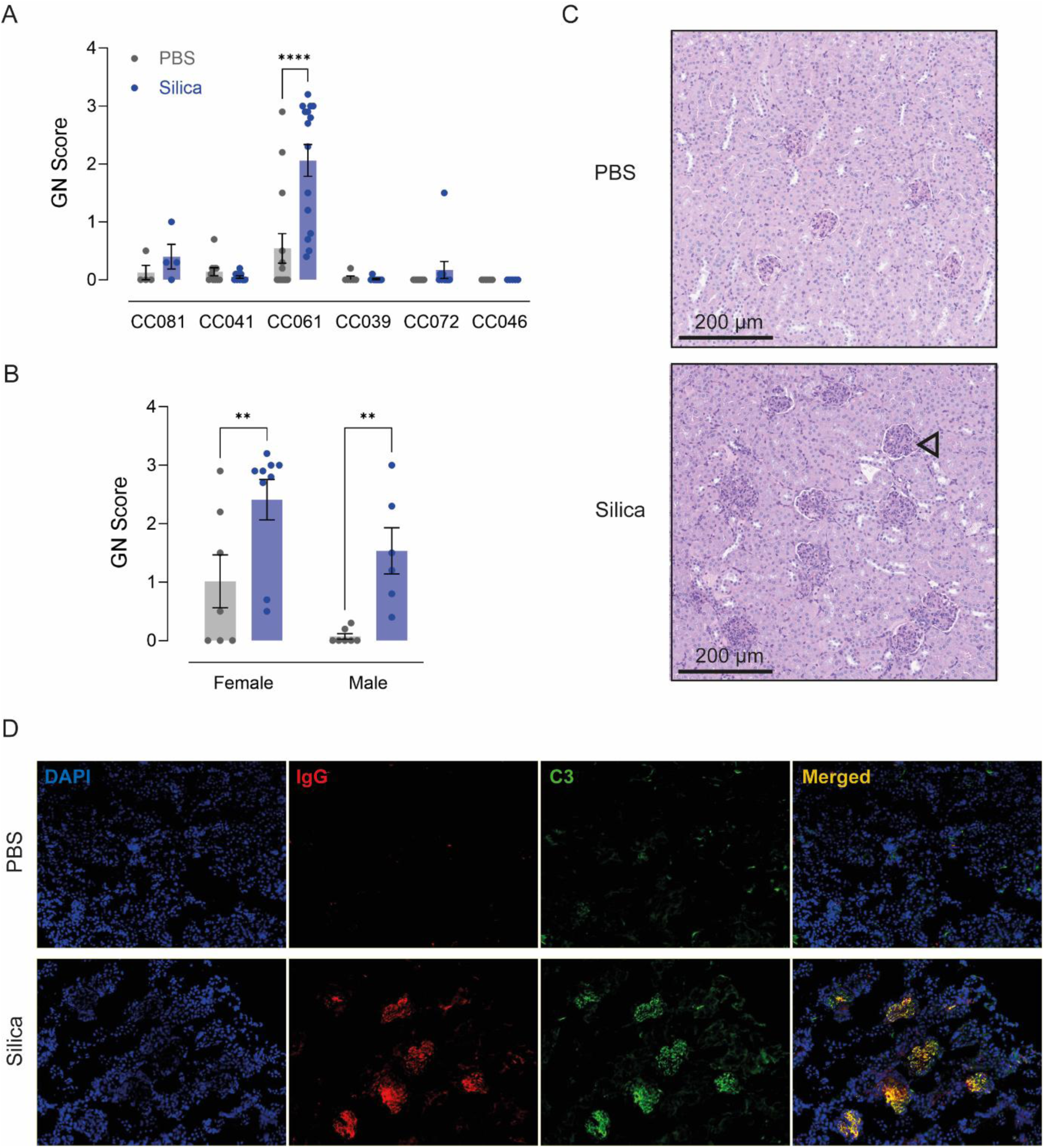
Silica exposure mediates glomerulonephritis in CC061 mice. (A) GN scores of silica- and PBS-exposed CC mice. (B) Sex comparison between GN scores of both PBS- and silica-exposed CC061 mice. Comparisons were performed using Two-Way Anova with Šídák’s correction for multiple testing. *p<0.05, **p<0.01, ***p<0.001, ****p<0.0001. (C) PAS-stained kidney tissue of PBS- and silica-exposed CC061, showing representative glomeruli. (D) Immunofluorescent staining of nuclei (DAPI), IgG, and C3 in kidney sections of PBS and silica-exposed CC061 mice.

### 3.5. Autoantibody profiles in silica-exposed mice are strain specific and exacerbated by susceptibility to sub-clinical autoimmunity

Development of idiopathic lupus or RA in humans is linked to the presence of disease specific autoantibodies [71] which are recapitulated in lupus [36] and RA [72] following occupational silica dust exposure. Autoantibody responses against selected disease-related autoantigens in the 6 silica-exposed CC RI strains revealed both strain and disease specific autoantibody profiles (Figure 5). Specifically, inflammatory arthritis in the CC039 was associated with RA linked anti-citrullinated protein antibodies (anti-CCP3) as well as ANA detected by IIF and ELISA, and anti-ENA6. In contrast, lupus-like disease in CC061 mice was linked to increased ANA (ELISA) and anti-chromatin but not anti-CCP3 or anti-ENA6. Both CC039 and CC061 had elevated serum IgG in response to silica exposure. The other four strains also exhibited strain-dependent autoantibody profiles including CC041 and CC081 mice which failed to develop silica-induced responses apart from a modest anti-ENA6 response in the CC081. CC046 mice were only positive for anti-ENA6 while CC072 mice had elevations in ANA (ELISA), anti-ENA6 and RF. PBS-exposed CC072 mice also had the highest mean values for ANA (IF), ANA (ELISA), anti-chromatin, and anti-CCP3 as well as significantly higher IgG titers compared to PBS mice from all other strains, with p <0.0001 for all comparisons. Silica-exposed CC072 mice also had significantly higher IgG titers compared to silica-exposed mice from other strains, with the exception of CC039. In addition to disease specific autoantibodies, silica exposure has been shown to elicit antibody responses against a diverse array of autoantigens in lupus-prone NZBWF1 mice [73].

**Figure 5:**
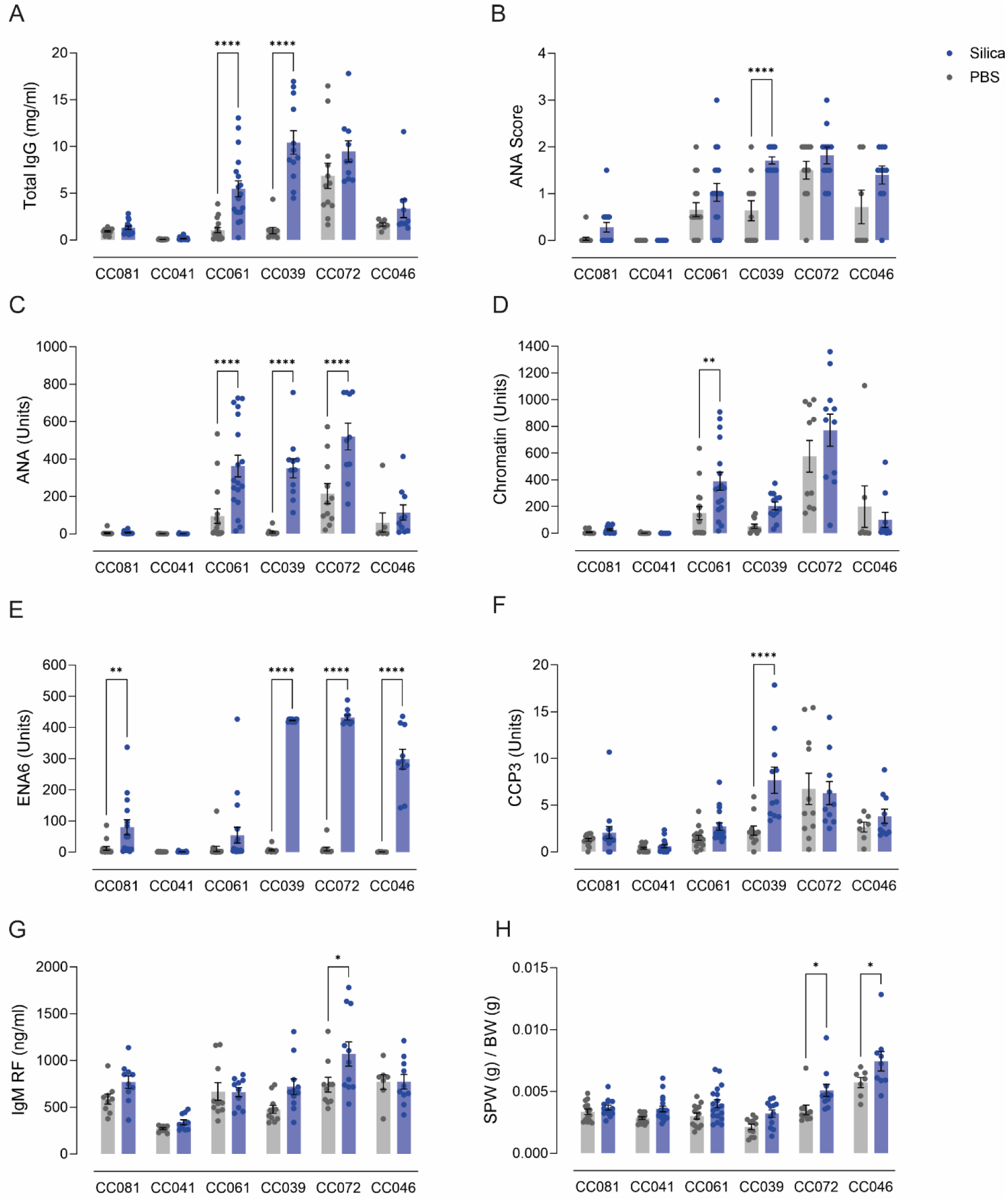
Silica-induced systemic autoimmunity in CC mice. Bar plots of IgG, autoantibodies and spleen weight. (A) Total IgG (mg/ml), (B) ANA IIF scores, (C) ANA (Units), (D) Chromatin (Units), (E) ENA6 (Units), (F) CCP3 (Units), (G) IgM RF (ng/ml), (H) SPW/BW. Comparisons between PBS-silica were performed using Two-Way ANOVA with Šídák’s correction for multiple testing. *p<0.05.

To determine whether differences in susceptibility to subclinical autoimmunity impact broader systemic immune activation, IgG autoantibody profiles were assessed using an autoantigen array. Principal component analysis (PCA) showed that the degree of separation between silica- and PBS-treated mice differed across strains (Supplementary Figures 2-4; (2) CC041 & CC081, (3) CC061 & CC039, (4) CC072 & CC046). The most striking separation was observed in CC039, where PBS and silica groups formed clearly distinct clusters, indicating a substantial silica-induced shift in the autoantibody landscape. CC046 and CC072 also demonstrated partial separation, suggesting treatment-related changes in these strains, although some overlap remained between groups. In contrast, CC041, CC081, and CC061 showed minimal or no distinct clustering, reflecting either a weaker or more heterogeneous response to silica exposure. These PCA findings were supported by heatmap analyses (Supplementary Figures 2-4), which revealed marked increases in both the number and intensity of IgG autoantibody responses in silica-exposed mice. Although each strain displayed a unique baseline autoantibody profile, silica exposure led to a substantial expansion in both the number and intensity of IgG responses. The number of significantly elevated autoantibodies ranged widely across strains. CC039 and CC061 exhibited the most extensive responses, with 90 and 79 upregulated targets, respectively, whereas CC041 and CC081 showed markedly smaller responses, with only 0 and 8 autoantibodies elevated, respectively. Strains with a higher burden of subclinical autoimmunity in the naïve state—such as CC039, CC061, and CC072—showed more diverse and expansive silica-induced autoantibody responses.

A striking sex effect was observed in CC046, where female mice exhibited a broader and more intense autoantibody response than males following silica exposure. This difference was clearly evident in the heatmap, where silica-exposed females clustered separately and showed denser autoantibody reactivity, suggesting that sex may interact with genetic background to modulate immune responses in a strain-dependent manner. No clear sex-related differences were observed in the other strains.

The number of significantly upregulated IgG autoantibodies varied widely by strain (Table 2), ranging from 0 in CC041 to 90 in CC039. CC046 and CC081 exhibited modest responses (11 and 8, respectively), while CC072 and CC061 displayed intermediate-to-strong profiles, with 69 and 79 upregulated targets. These differences broadly correlated with each strain’s baseline level of subclinical autoimmunity, with CC041 and CC081 showing the weakest induction and CC039, CC072, and CC061 exhibiting robust responses following silica exposure.

**Table 2:** IgG autoantibody responses elevated in silica exposed mice.

| CC RI Strain | IgG |
| --- | --- |
| CC041 | 0 |
| CC081 | 8 |
| CC046 | 11 |
| CC072 | 69 |
| CC039 | 90 |
| CC061 | 79 |

Among the upregulated autoantibodies, several targets were shared across multiple strains, particularly among those with stronger autoimmune phenotypes. A notable overlap was observed between CC039, CC046, CC061, and CC072, which all developed IgG autoantibodies against alpha fodrin, thyroglobulin, islet antigen-2 (IA-2), and interleukin-6 (IL-6). These targets are consistent with systemic autoimmune processes and are often associated with diseases such as lupus and autoimmune thyroiditis. Additionally, several nuclear and RNA-associated proteins—such as U1-snRNP, Sm/RNP complexes, and histones—were commonly upregulated among the high-responder strains, suggesting a strong activation of nuclear antigen-specific immunity. While the overlap among these strains was substantial, there were no IgG autoantibody targets shared across all six strains. In contrast to the shared responses, each strain also exhibited a unique set of autoantibodies that were selectively upregulated by silica, not detected in other strains. CC039 displayed the most diverse set of unique responses, including vimentin, SmD2, nucleolin, and collagens II, III, and IV, which may suggest enhanced tissue remodeling or joint involvement. CC072 also exhibited distinct autoantibodies such as GP2, dsDNA, and genomic DNA, along with U1-snRNP A, indicating a lupus-like humoral signature. CC061 uniquely upregulated several mitochondrial and cytoplasmic targets, including PR3, BPI, La/SS-B, and cytochrome C, as well as OGDC-E2, consistent with an innate immune activation profile. CC046 showed selective responses to U1-snRNP A, Sm/RNP, and CD40, potentially pointing to enhanced RNA-processing antigen presentation and immune co-stimulation. CC081 exhibited a more limited but distinct profile, including IFN-γ, lysozyme, and complement C1q.

### 3.6. Silica exposure induces strain-dependent cytokine/chemokine profiles

Silica exposure also elicited strain specific cytokine and chemokine profiles in several of the strains (Figure 6). CC072 mice, which were positive for the greatest number of inflammatory markers in naïve mice (Table 1), also had the greatest change in silica-induced inflammatory marker response with increases in IL-2, IFN-γ, MIG (Cxcl9) and MIP-1β (Ccl4). CC039 mice were the only other strain with sub-clinical autoimmunity to show a silica mediated change in inflammatory markers with an increase in G-CSF. In contrast, the two strains without sub-clinical inflammatory markers had increases in IL-17 and RANTES (CC081) and G-CSF (CC041) in silica exposed mice compared to corresponding controls.

**Figure 6:**
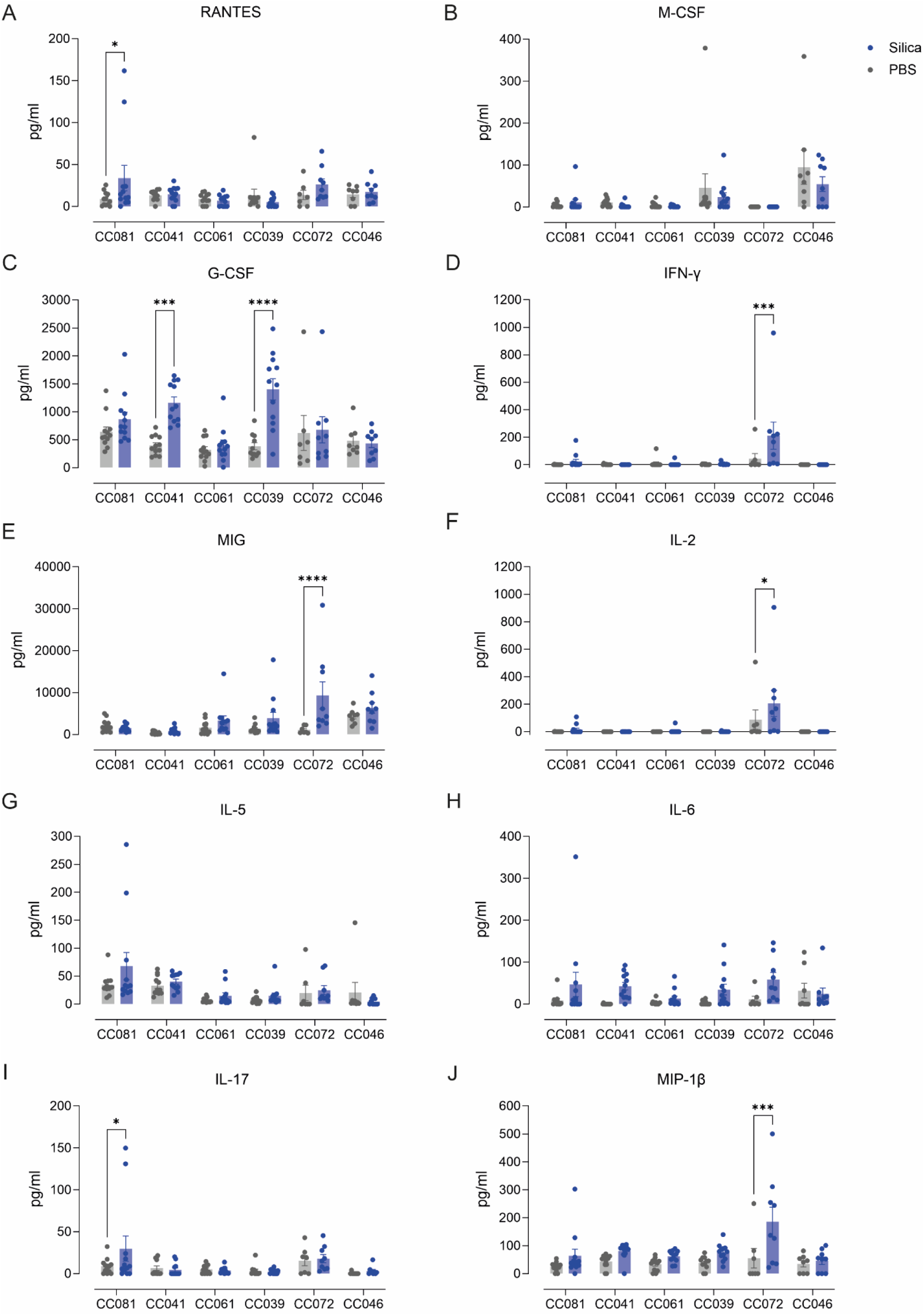
Cytokine/chemokine profiles in serum. Bar plots (pg/ml) of (A) RANTES, (B) M-CSF, (C) G-CSF, (D) IFN-γ, (E) MIG, (F) IL-2, (G) IL-5, (H) IL-6, (I) IL-17, (J) MIP-1β. Comparisons were performed using Two-Way Anova with Šídák’s correction for multiple testing.

### 3.7. Severity of silica-induced lung inflammation is strain-dependent and linked to development of autoimmunity

The mucosal origins hypothesis identifies the lungs as a major location in the development of RA [35], and other autoimmune diseases particularly following inhalant exposures such as silica dust [2]. Silica exposure in mice results in pulmonary inflammation characterized by alveolar, peribronchial, and perivascular inflammatory cell infiltrates, and tracheobronchial lymph node (TBLN) hypertrophy [34, 74]. While these initial inflammatory responses may vary depending on mouse strain [75], it remains unclear whether any of these silica-induced pulmonary inflammation biomarkers require pre-existing sub-clinical autoimmunity and whether they are linked to development of pathological disease. To determine the importance of susceptibility to sub-clinical autoimmunity in the development of silica-induced pulmonary pathology we examined a number of features in the lungs and draining lymph nodes of all 6 CC strains following silica exposure including alveolitis, perivasculitis and peribronchitis, lung granuloma (silica-containing nodules surrounded by inflammatory cells), fibrosis, proteinosis, enlarged TBLN, and formation of silicotic nodules in TBLN (Figure 7A and B.

**Figure 7:**
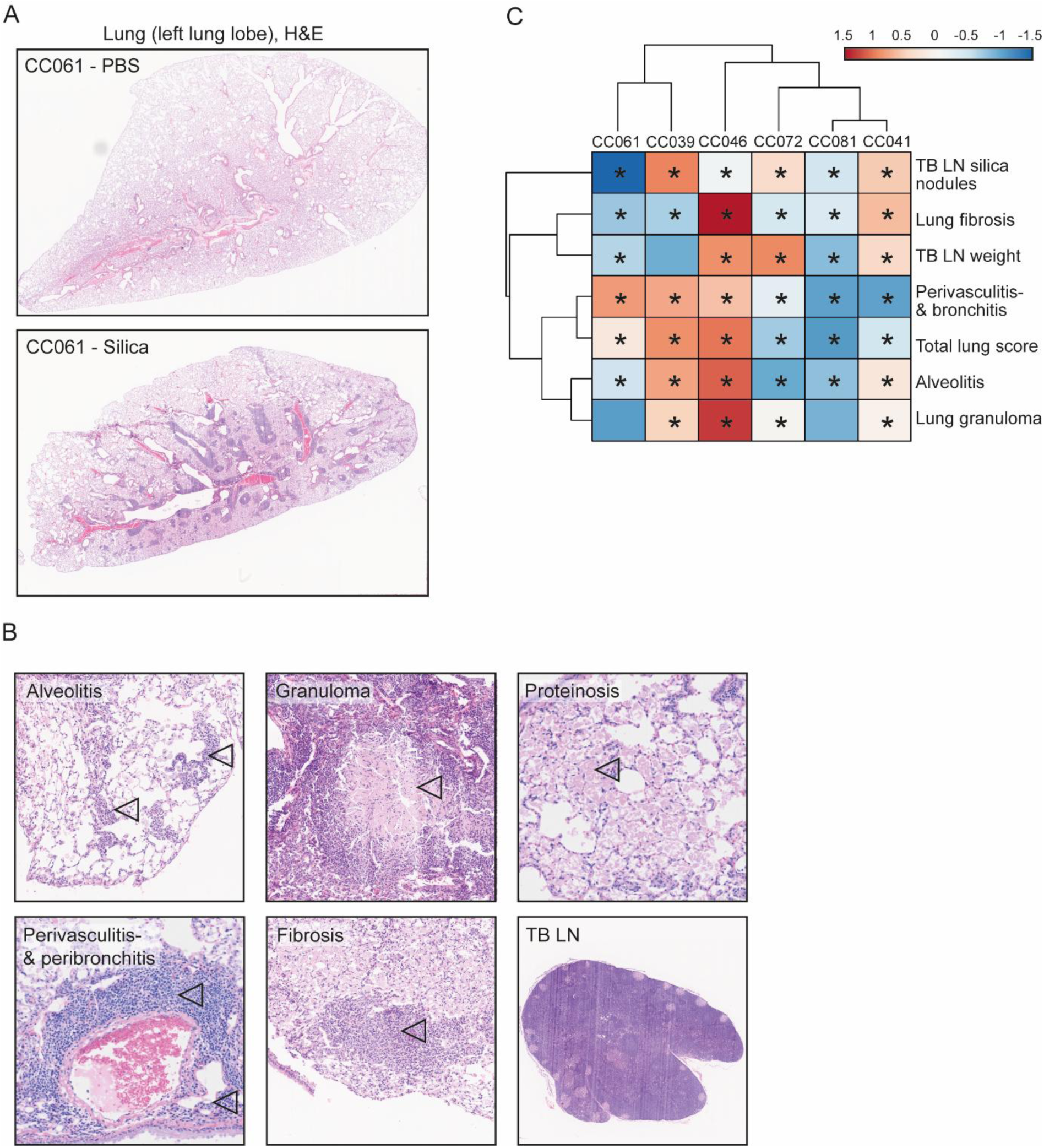
Silica-induced pulmonary inflammation biomarkers in CC RI mice. (A) Representative H&E-stained lung tissue from a silica-exposed and a PBS-exposed CC061 mouse, chosen for its moderate development of discussed silica-induced features. (B) Histological figures showing examples from all the assessed features (alveolitis, perivasculitis & peribronchitis, granuloma, fibrosis, proteinosis and enlarged TB LN). (C) Heatmap with hierarchical co-clustering of lung inflammation and silicosis biomarkers in silica-exposed mice. *p<0.05, indicates significant difference with PBS-exposed mice.

A heatmap (Figure 7C), comparing these features including co-clustering based on strain and lung pathology biomarkers, showed significant strain dependent differences which are represented numerically in Supplementary Figure 1. Significantly, CC RI strains that developed the most pronounced autoimmune responses upon silica exposure (CC039, CC061, CC046, CC072), were observed to have significantly more perivasculitis and peribronchitis inflammatory infiltrates than the strains lacking susceptibility to sub-clinical autoimmunity (CC081, CC041) (Supplementary Figure 1B). Additionally, the strains that developed disease specific pathology (arthritis in CC039 and nephritis CC061) clustered separately from the other strains (Figure 7C). Although naive CC046 and CC072 mice showed evidence of sub-clinical autoimmunity (Figure 1, Table 1), their pulmonary pathology indices clustered more closely with CC081 and CC041. In particular, the total lung scores for CC072 were more consistent with CC081 and CC041 (Supplementary Figure 1C) even though CC072 exhibited the most obvious evidence of sub-clinical features in naïve mice (Figures 1 and 5). In contrast, CC046 developed significant pulmonary inflammation but its silica-induced autoantibody response was limited (Figure 5). Although these findings show that silica-induced pulmonary inflammation is a common feature of the 6 CC strains studied, they also reveal that the severity of silica-induced perivasculitis and peribronchitis is the pulmonary response that best reflects the positive relationship between the susceptibility to sub-clinical autoimmunity and the development of more severe features of systemic autoimmunity.

## 4. Discussion

Crystalline silica exposure, a common occupational exposure, has been associated with several systemic autoimmune diseases [28, 76]. In this study, we used the Collaborative Cross recombinant inbred (CC RI) mouse lines to investigate if silica exposure in genetically related mice can lead to a spectrum of autoimmune pathologies that mimic those found in occupationally exposed humans. Our studies reached several important conclusions. First, we determined that CC RI strains exhibit a spectrum of sub-clinical autoimmunity based on autoantibodies and inflammatory markers. Second, we confirmed the connection between pre-existing subclinical autoimmunity and the onset and severity of silica-induced systemic autoimmunity. Finally, we revealed that silica exposure of CC RI strains leads to the development of distinctly different autoimmune diseases phenotypes. Importantly, our findings provide the first direct evidence that silica exposure can trigger inflammatory arthritis in mice, supporting the association between silica exposure and rheumatoid arthritis previously made based on epidemiological evidence [28, 45, 77, 78]. Furthermore, we confirm that silica exposure can exacerbate lupus-like disease, including glomerulonephritis. Using a comprehensive autoantigen array, we further demonstrated that silica exposure amplifies systemic autoantibody responses in a highly strain-specific manner. The number and specificity of IgG autoantibodies varied substantially across CC strains, with high-responder strains such as CC039, CC061, and CC072 exhibiting broad and diverse autoantibody profiles, while others like CC041 remained largely unresponsive. Notably, several autoantigens—including alpha fodrin, IA-2, thyroglobulin, and IL-6—were commonly elevated in susceptible strains, suggesting shared immunological pathways underlying silica-induced autoimmunity. These findings provide molecular-level evidence that supports a mechanistic link between genetic predisposition and silica-triggered systemic autoimmunity. Additionally, our findings strengthen the premise that susceptibility to sub-clinical autoimmunity is a significant risk factor in the development of silica-induced autoimmune disease.

Analysis of ANA and inflammatory cytokines and chemokines in serum from 61 naïve CC RI lines revealed a broad spectrum of immune activity, ranging from strains with no detectable autoimmune or inflammatory markers—resembling immunologically “healthy” individuals—to strains exhibiting clear serological signs of subclinical autoimmunity. These findings demonstrate that the genetic diversity of the CC RI panel naturally gives rise to varying baseline immune profiles, consistent with what is observed in human populations. In humans, subclinical autoimmunity is increasingly recognized as a precursor state to systemic autoimmune disease, including idiopathic conditions such as lupus and rheumatoid arthritis [79]. The identification of CC strains with distinct immune signatures, including those that are ANA-negative yet positive for inflammatory mediators associated with autoimmunity, mirrors similar immune endotypes described in human patients [9]. Interestingly, strains that were ANA-negative but expressed autoimmunity-associated cytokines also overlapped with those reported in humans to be at increased risk of progression to clinical disease despite negative ANA status [9]. However, it is important to recognize that the current dataset provides only a single time-point “snapshot” of immune activity in these strains. Unlike longitudinal studies in humans that can track the progression from subclinical to overt disease, our cross-sectional design does not capture how these immune profiles may evolve over time within individual mice.

Significantly, manifestations of systemic autoimmune disease, specifically glomerulonephritis and inflammatory arthritis, and the presence of autoantibodies, was observed in those strains selected as positive for sub-clinical autoimmunity. These findings strengthen the premise that susceptibility to preclinical autoimmunity is a significant risk factor in the development of silica-induced autoimmune disease. Silica-induced pulmonary pathology in the CC RI lines ranged from mild to severe, with variable degrees of inflammation in peribronchial, perivascular, and alveolar regions, and variable degrees of lung fibrosis and lung granuloma. These observations are in agreement with prior studies on a selection of mouse strains, which have recorded significant variability in both the severity and types of lung lesions post silica particle inhalation [75]. Similarly, this variability in response was also detected in silica-exposed diversity outbred (DO) mice [34]. Whether the extent of lung inflammation or silicosis in humans is associated with the development of autoimmunity upon silica exposure, and whether more extensive lung inflammation and autoimmunity are driven by the same genetic background, is unclear. Based on our data, it appears that perivascular and peribronchiolar inflammation is more apparent in those strains (CC039, CC061, CC046 and CC072) that develop higher levels of autoantibodies, spontaneously or upon silica exposure.

The strains that do develop autoimmunity as evidenced by autoantibody levels upon silica exposure, develop different autoantibody profiles. CC072 and CC039 develop a very clear anti-ENA6 response upon silica exposure, while CC061 develop an anti-chromatin response upon silica exposure. The ENA responses upon silica appeared to be less variable compared to the ANA and anti-chromatin responses, which were also present in some of the PBS mice. These findings are similar to findings with the DO mice, in which the ENA response of ANAs was a clear silica-associated response, as PBS mice that were ANA positive generally lacked the specific ENA reactivity [34]. Whether ENA, such as anti-Smith (Sm) and anti-Ribonucleoprotein (RNP) are involved in pathogenesis, is unclear and warrants further investigation.

Interestingly, we found that silica exposure in CC039 mice is associated with the development of inflammatory arthritis. Silica-exposed CC039 mice develop significant synovial inflammation and bone erosions in ankle joints, that is similar, but less severe, compared to the pathology observed in collagen-antibody induced arthritis (CAIA). The development of arthritis in CC039 mice exhibited a pronounced female bias, consistent with the well-documented female predominance in autoimmune diseases. This pattern has been linked to X-linked factors such as Xist, which has been shown in mouse models to play a role in immune regulation and autoimmunity [80]. Whether the inflammatory arthritis observed in these mice is the result of an autoimmune reaction, and could therefore have overlap in pathogenesis with RA in humans, is not clear yet. However, the significant silica-induced autoantibody responses in CC039 mice suggest that the arthritis could be a result of autoimmunity. Furthermore, it is unclear whether the CC039 mice spontaneously develop inflammatory arthritis, or whether this is a silica-specific response. Mice were harvested 12- and 20 weeks post-exposure, which aligns with 20-32 weeks of age. In another mouse strain that spontaneously develops inflammatory arthritis, the BXD2/TyJ mouse, about 50% of female mice develop arthritis by 34 weeks of age [81]. The fact that none of the PBS-exposed mice were positive for inflammatory arthritis, suggests a slower development of disease in the CC039 mice, or the absence of the spontaneous development of the disease.

In human arthritis, antibodies against cyclic citrullinated proteins (ACPA/CCP) and rheumatoid factor (RF, IgM) are hallmarks of disease, and have diagnostic relevance [82]. Notably, CC039 mice showed the strongest development of ACPA/CCP of the included CC RI strain, and it was the only strain with significantly higher titers of CCP in silica-exposed mice compared to PBS mice. These findings confirm similarities between human RA and the phenotype observed in CC039 mice. Whether CCP antibodies play a causal role in the development of disease in these mice, is unclear, but the absence of a significant correlation between synovitis scores and CCP suggests that CCP levels are not predictive for arthritis in CC039 mice, such as is the case in RA patients. RF IgM on the other hand was not significantly increased in silica-exposed mice, and was not higher in CC039 mice compared to other CC RI strains.

As none of the other CC RI lines, and none of the CC RI founder strains, developed inflammatory arthritis, it becomes clear that a specific combination of genetic traits in combination with silica exposure, is what drives the disease in these mice. This phenomenon has been observed previously with the BXD RI lines, derived from the C57BL/6J and DBA/2 parental strains. Neither C57BL/6J nor DBA/2 mice spontaneously develop arthritis [81], and both are resistant to collagen-induced arthritis [83, 84]. However, one BXD RI strain, the BXD2 mouse, develops severe generalized autoimmune disease with inflammatory arthritis [81], showing that adventitious mixing of genetic variants from non-autoimmune strains can produce an autoimmune genotype.

Apart from the inflammatory arthritis in CC039 mice, we found that CC061 develops a lupus-like phenotype upon silica exposure, presented with the development of glomerulonephritis (GN). Moreover, also a portion of the female PBS-exposed mice develop GN, indicating that these mice likely spontaneously develop GN over time, and that 20-22 weeks of age is too early to consistently observe spontaneous GN. Although lupus is not a major characteristic of any of the CC RI founder strains, lupus-predisposing loci have previously been described in three, C57BL6/J, 129S1/SvImJ, and NOD/ShiLtJ [85, 86]. The CC061 has inherited its MHC class II genes from the 129S1/SvImJ mouse (chromosome 17). Moreover, CC061 has also inherited the distal part of Chromosome 1 from the 129S1/SvImJ mouse, which has been proven sufficient to mediate the loss of tolerance to nuclear antigens in a C57BL/6 congenic strain carrying the 129S1/SvImJ Chromosome 1 segment [87, 88]. While neither the pure 129S1/SvImJ nor the C57BL/6 mouse strains were observed to develop autoimmunity spontaneously, a lupus-like condition has been observed in hybrid strains resulting from a cross between 129S1/SvImJ and C57BL/6 [89, 90]. This indicates that the hybrid mice’s tendency towards this condition might stem from the combination of specific alleles inherited from both the 129S1/SvImJ and C57BL/6 parent strains, which could explain the observation of lupus-like disease in the CC061 mice. A sex effect was observed in the development of GN in CC061 mice, in favor of more GN development in female mice. In addition, autoantibody levels were higher in female CC061 mice. Although these findings are in line with observations in humans, where a higher prevalence is observed in women [23].

Lastly, our findings also argue that the genetic diversity of the CC RI panel provides a powerful model with which to study the relationships between genetics, sub-clinical autoimmunity, and xenobiotic/environmental exposures in the development and severity of autoimmune disease. The observation that silica triggers differential responses across various CC RI strains, which all inherit genetic material from the same eight founder strains [55], presents a unique opportunity to study the genetic background involved in silica-mediated autoimmunity. Other recombinant inbred lines, such as the BXD and NZM, have contributed to the genetic exploration of autoimmunity but are limited by their genotypes and/or commercial availability [51, 54, 81] The ongoing development of CC RI strains, alongside advancements in study protocols [91, 92], informatics resources [93, 94], QTL mapping software [95, 96], and founder strain-specific SNP arrays [97], highlights the CC RI panel’s evolving utility for genetic research. Furthermore, the increased number of intercross generations within the CC strains is expected to facilitate more rapid narrowing of intervals compared to traditional F2 intercrosses or backcrosses. However, conducting genetic analyses using quantitative trait loci (QTL) analyses, would necessitate more than 30 CC strains [91, 98]. These requirements exceed the capabilities of the current study, which primarily serves as a proof of principle experiment. Thus, future studies should expand the number of included CC RI strains to effectively use QTL analyses to investigate genetics of silica-induced autoimmunity.

## 5. Conclusions

In conclusion, this study affirms our hypothesis that susceptibility to subclinical autoimmunity significantly influences the development and severity of silica-induced systemic autoimmunity. By leveraging the genetic diversity inherent in the Collaborative Cross strains, we demonstrate a clear correlation between the presence of autoantibodies and inflammatory markers before exposure and the subsequent development of several systemic autoimmune diseases, including inflammatory arthritis and lupus-like disease with and without glomerulonephritis following silica exposure. Moreover, our research underscores the potential of using genetically diverse animal models to predict disease susceptibility and establishes the CC RI strains as an important tool for studying environment-induced autoimmunity and gene-environment interactions in the induction of autoimmunity.

## Declarations

### Ethics approval and consent to participate

All animal procedures were approved by TSRI Institutional Animal Care and Use Committee (IACUC) (protocol# 08-0150).

### Competing interests

The authors declare no competing interests.

### Funding sources

This research was funded by NIH grants via the National Institute of Environmental Health Science (NIEHS) and National Institute of Allergy and Infectious Diseases (NIAID) under Award numbers R01ES029581, R21ES031454, and R21AI174022 to KMP. LMFJ was supported by FWO (Research Foundation - Flanders) grants V420121N and V412323N. PHMH was supported by KU Leuven [Internal Funding C2 (C24/18/085)].

### CRediT authorship statement

**Lisa M.F. Janssen:** *Conceptualization*, *Data curation, Formal analysis*, *Funding acquisition*, Investigation, Methodology, Project administration, Resources, Software, Supervision, Validation, Visualization, *Writing – original draft, Writing – review and editing*

**Caroline de Ocampo:** Conceptualization, *Data curation*, Formal analysis, Funding acquisition, Investigation, Methodology, Project administration, Resources, Software, Supervision, Validation, Visualization, Writing – original draft, *Writing – review and editing*

**Dwight H. Kono:** *Conceptualization*, *Data curation*, Formal analysis, Funding acquisition, Investigation, Methodology, Project administration, Resources, Software, Supervision, Validation, Visualization, Writing – original draft, *Writing – review and editing*

**Peter H.M. Hoet:** Conceptualization, Data curation, Formal analysis, *Funding acquisition*, Investigation, Methodology, Project administration, Resources, Software, Supervision, Validation, Visualization, Writing – original draft, *Writing – review and editing*

**K. Michael Pollard:** *Conceptualization*, Data curation, Formal analysis, *Funding acquisition*, Investigation, Methodology, *Project administration*, Resources, Software, *Supervision*, Validation, Visualization, Writing – original draft, *Writing – review and editing*

**Jessica M. Mayeux:** *Conceptualization*, *Data curation*, Formal analysis, Funding acquisition, Investigation, Methodology, Project administration, Resources, Software, *Supervision*, Validation, Visualization, Writing – original draft, *Writing – review and editing*

## Data availability

The datasets used and/or analyzed during the current study are available from the corresponding author on reasonable request.

## Notes

### Competing Interest Statement

The authors have declared no competing interest.

