## Supplementary Materials for "Crystalline silica exposure induces multiple systemic autoimmune phenotypes including inflammatory arthritis and nephritis in Collaborative Cross mice with differing sub-clinical autoimmune profiles"

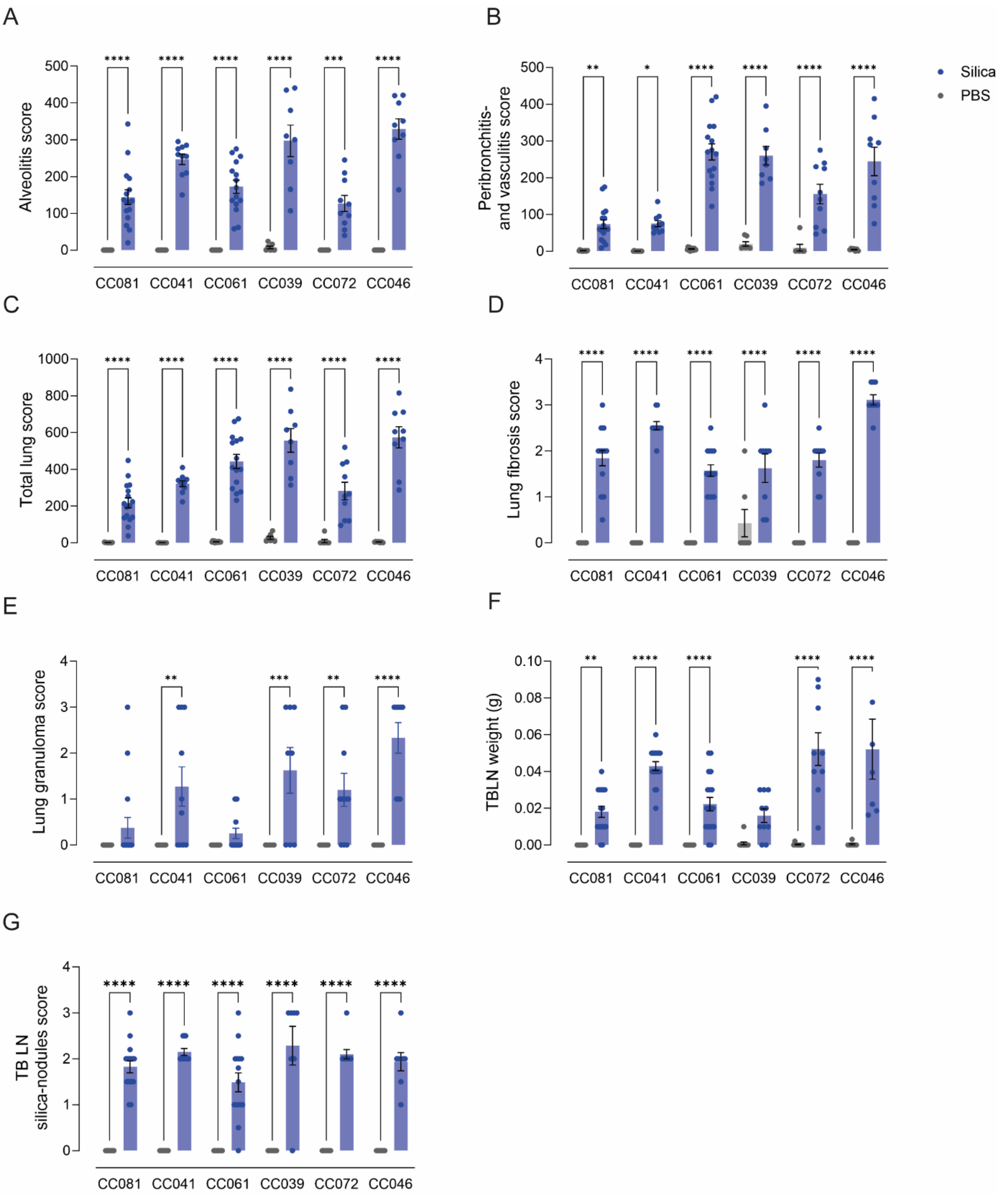


**Supplementary Figure 1: Pulmonary inflammation scores.** (A) Alveolitis score, (B) Peribronchitis- and perivasculitis score, (C) Total lung score, (D) Lung fibrosis score, (E) Lung granuloma score, (F) TB LN weight (g), (G) TB LN silica-nodules score. Comparisons were performed using Two-Way Anova with Šídák’s correction for multiple testing.

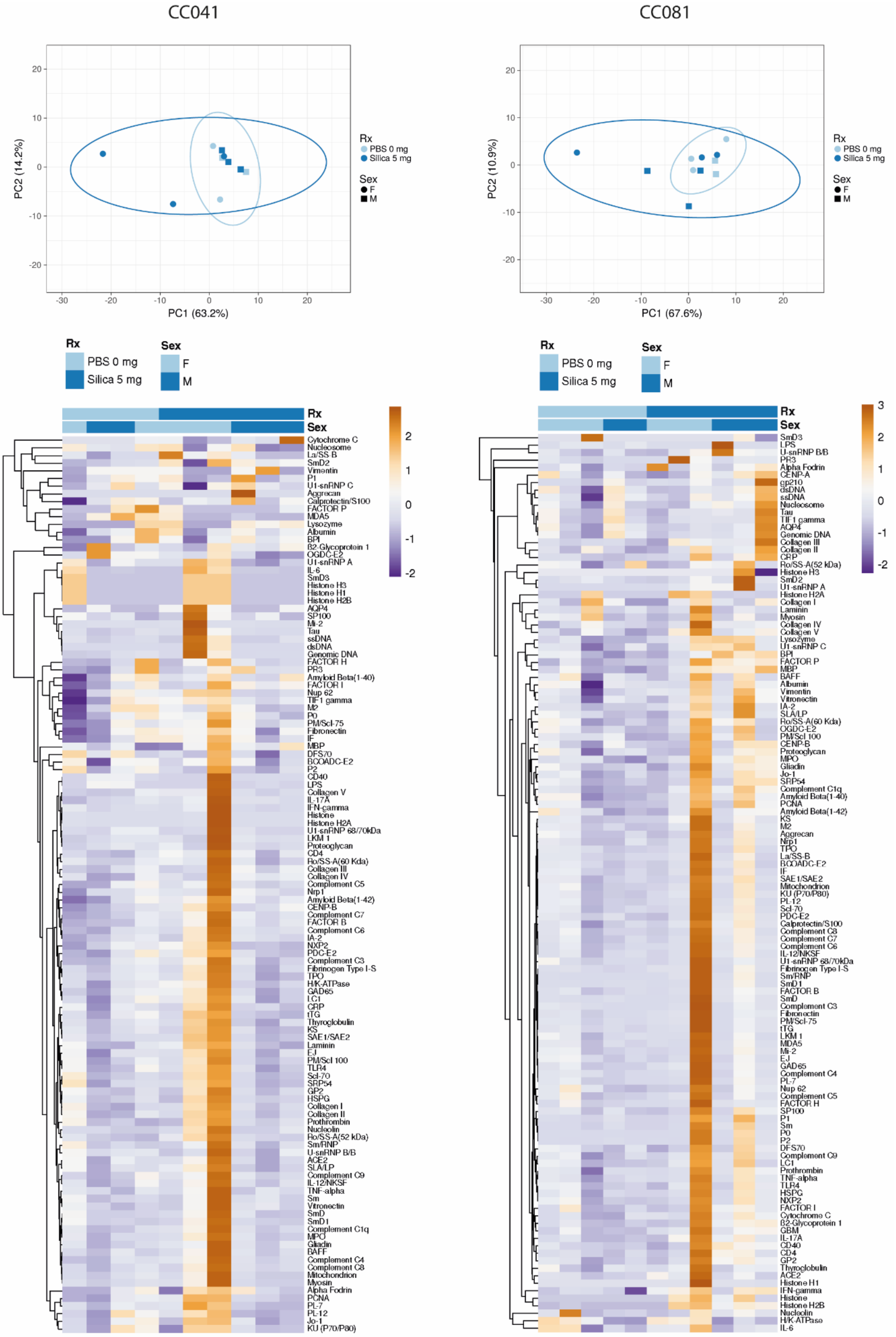


**Supplementary Figure 2:** PCA and heatmaps with clustering for autoantibodies of autoantigen array data for CC041 and CC081.

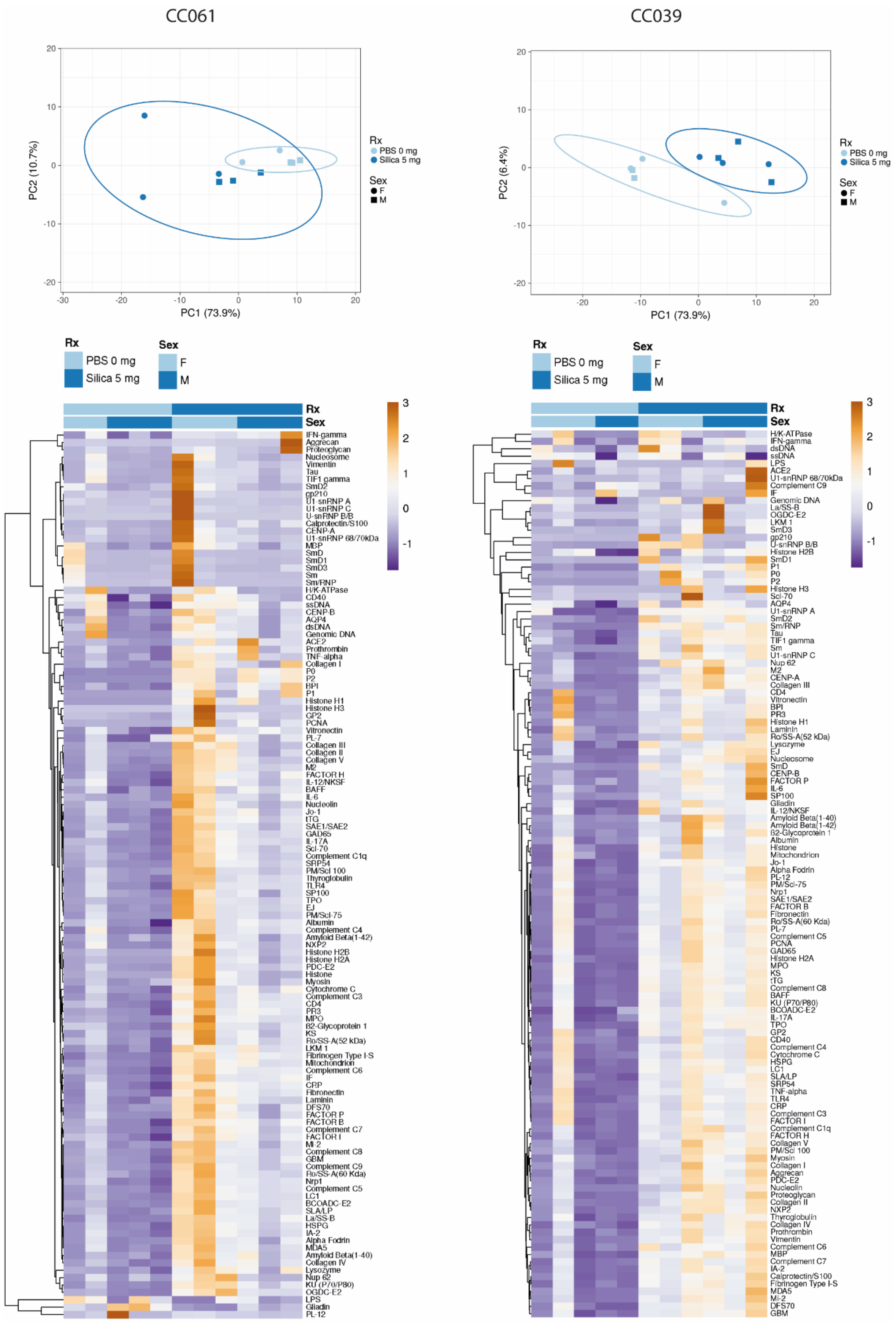


**Supplementary Figure 3:** PCA and heatmaps with clustering for autoantibodies of autoantigen array data for CC061 and CC039.

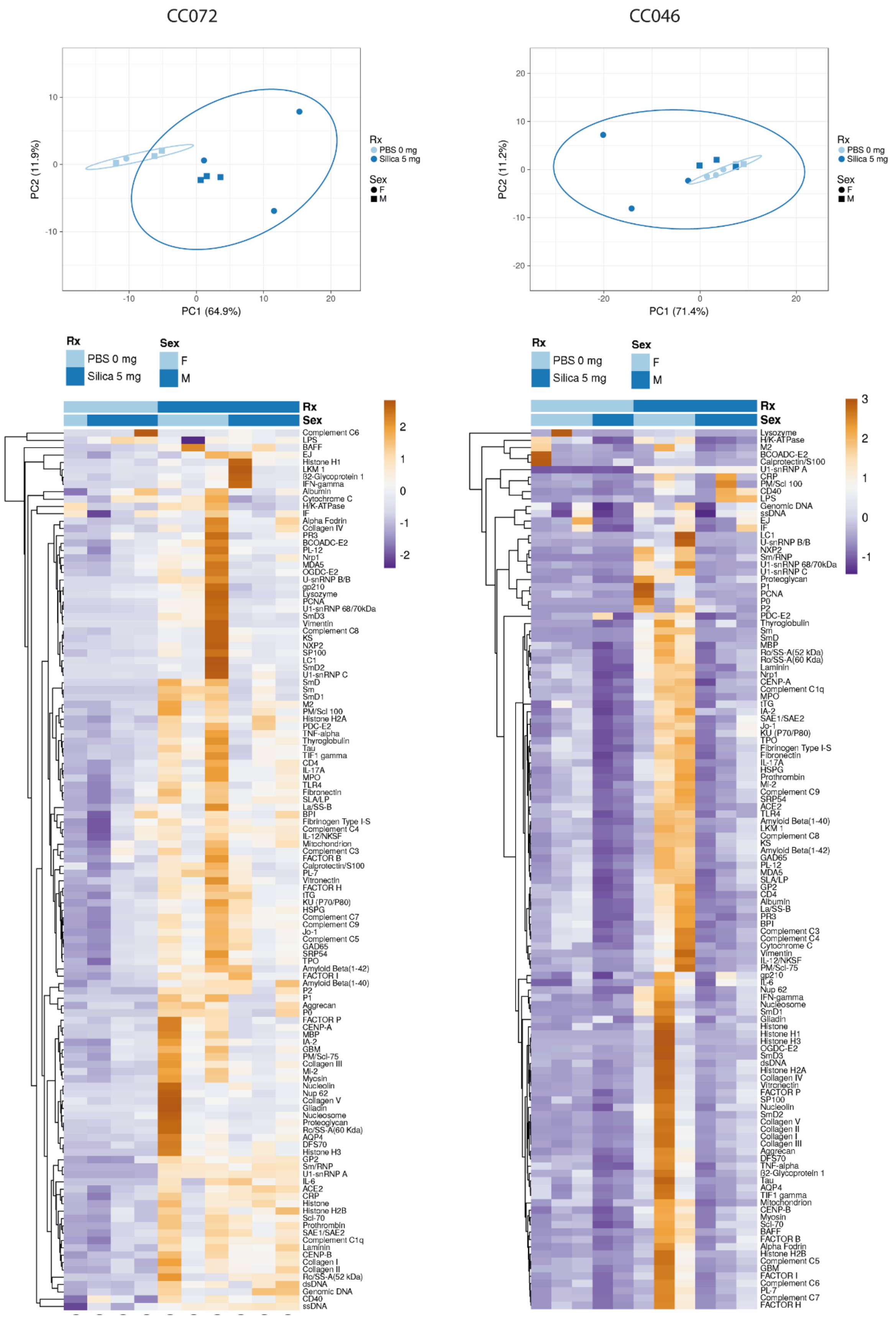


**Supplementary Figure 4:** PCA and heatmaps with clustering for autoantibodies of autoantigen array data for CC072 and CC046.
